# Latent mitotic vulnerability of AML cells induced by therapeutic agents

**DOI:** 10.1101/2024.06.27.600939

**Authors:** Ryuta Niikura, Tomohiro Yabushita, Shohei Yamamoto, Hiroaki Suzuki, Masamitsu Fukuyama, Shoji Hata, Susumu Goyama, Toshio Kitamura, Takumi Chinen, Daiju Kitagawa

## Abstract

Acute myeloid leukemia (AML) is a hematopoietic malignancy with a poor prognosis. Understanding the unidentified properties of AML cells is beneficial for the identification of novel therapeutic strategies for AML. In this study, we uncover the vulnerabilities of AML cells in mitosis when exposed to therapeutic agents. Through comparative analysis of large-scale data quantifying drug effects on cancer cell proliferation, the drug targeting the cell cycle and mitosis are predicted to possess high cytotoxicity against AML cell lines. Consistently, live-cell imaging with microwell devices demonstrates that clinical drugs targeting the cell cycle processes, such as idarubicin, pevonedistat and vincristine, potently induce mitotic cell death in AML cells. While these therapeutic agents also induce cell death through S/G2 phase arrest, the cytotoxic effects during mitosis are notably more pronounced. Furthermore, by employing additional inhibition of Chk1 to override the G2/M checkpoint, the AML cells stalled in the S/G2 phase prematurely enter mitosis, resulting in a significant increase in cell death. Collectively, these results unveiled the latent mitotic vulnerabilities of AML cells, providing a basis for developing novel therapeutic interventions.

## Introduction

Acute myeloid leukemia (AML) is a hematological malignancy characterized by abnormal proliferation and accumulation of immature myeloid cells^1,2^. The estimated 5-year survival rate for AML reported since 2020 is 31%, indicating a poor prognosis for this disease^3^. Because the response to the chemotherapy greatly impacts treatment outcomes against AML, the development of novel therapeutic agents with higher efficacy has been continuously pursued^4–6^. Accordingly, with an improved understanding of the genetic aberrations of AML, various molecular targeted therapies have been clinically applied since 2017^2,7–9^. However, due to the complexity of genetic aberrations in AML, selecting the optimal treatment remains challenging^9–11^. In addition, current molecular targeted therapies have yet to improve the treatment outcomes against poor prognostic factors^9–12^, such as complex chromosomal abnormality or ASXL1 mutation. Therefore, to advance AML treatment, it is crucial to investigate AML cell characteristics and vulnerabilities using methods beyond genetic aberration analysis, facilitating the development of innovative therapeutic strategies.

Comparing gene dependencies and drug responses to cell proliferation across different cell types is a key strategy for identifying therapeutic vulnerabilities specific to each cell type^13–15^. In line with this, a comparative approach that contrasts the responses to gene knockout and drug treatment between AML cells and other cell types could provide valuable insights to elucidate AML-specific vulnerabilities. To date, genome-wide CRISPR/Cas9 screening and drug response analyses of barcoded tumor cell pools have been conducted for various cancers, including AML^15,16^. These large-scale data have been registered in the Cancer Dependency Map (DepMap)^17,18^. Recent studies utilizing the CRISPR/Cas9 screening data of the DepMap project have presented AML subtype-specific drug targets^19^. Furthermore, it has been shown that, according to the drug sensitivity data from DepMap, decitabine, a DNA methyltransferase inhibitor used for AML treatment, exhibits higher sensitivity in AML cells compared to non-hematopoietic cancers^20^. Thus, leveraging large-scale data and tools including DepMap potentially leads to a comprehensive understanding of AML cell characteristics and contributes to the development of new therapeutic strategies targeting the specific vulnerabilities of AML. Live-cell imaging is a useful technique for a deeper understanding of the cellular responses to therapeutic agents^21–25^. For instance, detailed dissection of the mitotic processes has shown that decitabine treatment induces a high frequency of chromosome missegregation in AML cells associated with poor prognosis^20^. Thus, by using live-cell imaging, the analysis of long-term cellular responses to therapeutic agents can provide new insights into understanding the characteristics of AML cells. However, because most AML cells are suspension cells and easily flow in the medium, long-term tracking of individual cells is technically challenging.

In this study, utilizing the comprehensive dataset of drug sensitivity from the DepMap and an optimized microwell-base live-cell imaging, we revealed that therapeutic agents targeting the “cell cycle” and the “mitosis” are effective against AML cells. These therapeutic agents rapidly elicited mitotic cell death in AML cells. Thus, the mitotic vulnerability of AML cells provides a basis for proposing new treatment strategies targeting AML.

## Results

### The DepMap data predicts drugs targeting the “cell cycle” and “mitosis” to possess potentially high cytotoxicity against AML cell lines

We explored potential vulnerabilities of AML cells by comparing the drug sensitivities of AML cells to those of non-hematological malignancy (non-HM) cells (Fig. 1A). We first analyzed the drug sensitivity AUC (CTD^2) data of available cancer cell lines from DepMap, which suggested that certain drugs potentially exhibit higher cytotoxicity against AML cell lines compared to non-hematopoietic cancer cell lines (Fig. 1B, 545 drugs). Among them, we selected the top 27 drugs (Effect size ≤-3.0) for subsequent analysis. The molecular targets for these drugs were extracted from the Drug Repurposing Hub^26^ and the Cancer Therapeutics Response Portal v2^27^ (Fig. 1A, C). Then, we performed unbiased Gene Ontology (GO)^28^ analysis on these extracted drug targets (41 genes). The results suggested that AML cells tended to be sensitive to drugs targeting factors involved in biological processes (BP) related to cell cycle and mitosis, as well as cellular components (CC) associated with centromeric region, chromosomal region, and microtubules (Fig. 1D, 1E). On the other hand, AML cells do not appear to be sensitive to drugs targeting factors associated with other GO terms, such as response to organonitrogen compounds and the gamma-secretase complex (Fig. S1A-D). These results suggest that AML cells might be sensitive to drugs targeting cell cycle progression and mitotic phase.

**Figure 1.**
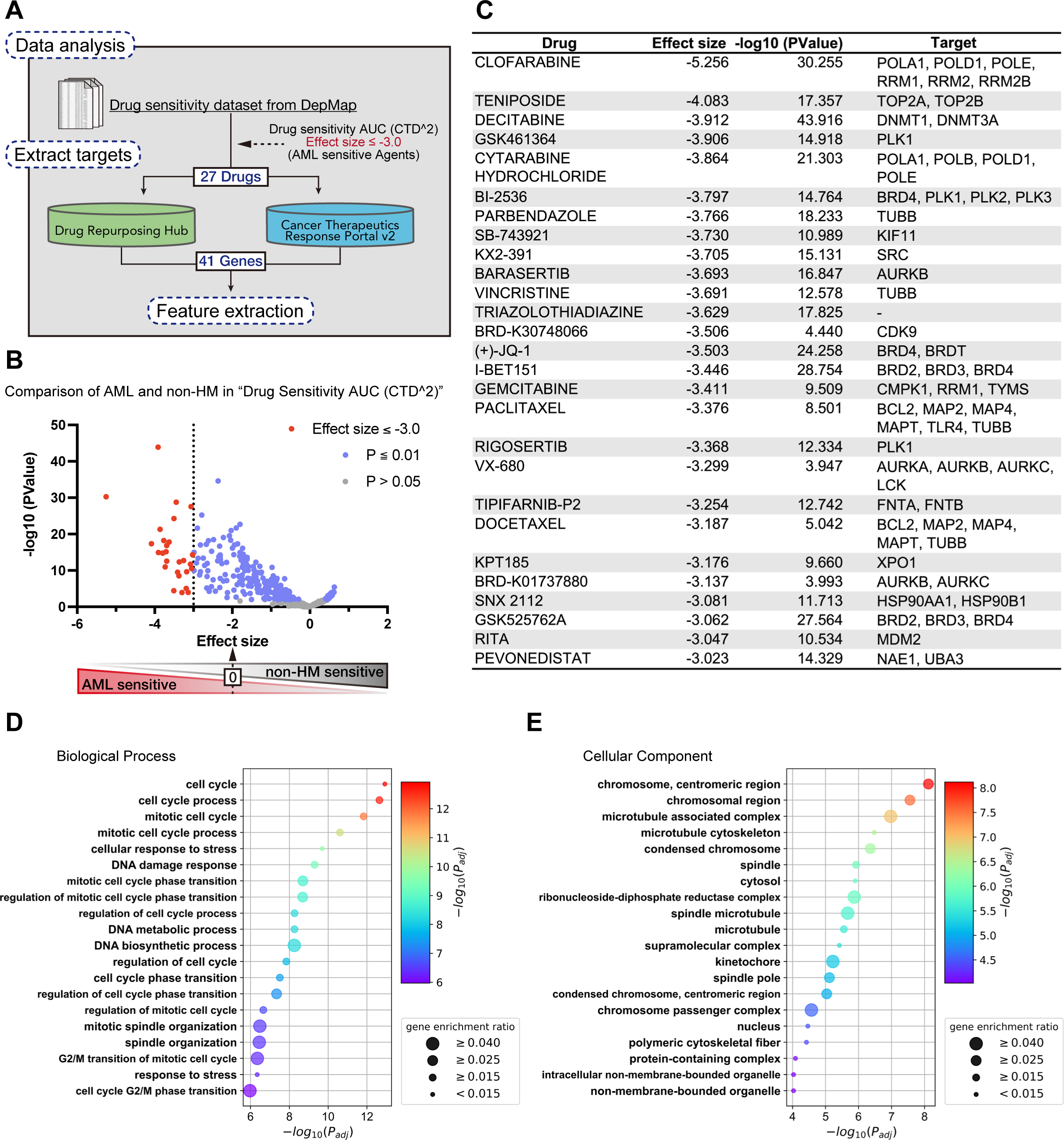
DepMap data predicts that drugs targeting the “cell cycle” and “mitosis” are highly effective against AML cells. (**A**) Scheme for exploring the potential vulnerabilities of AML cells by comparing drug sensitivity AUC (CTD^2) data from the DepMap. (**B**) Volcano plots comparing drug susceptibility in AML cells and non-hematopoietic malignancy (non-HM) cells. (**C**) Drugs and their targets predicted to be highly effective to AML cells based on the analysis shown in (A) and (B). (**D, E**) Gene Ontology (GO) enrichment analysis using g.Profiler was conducted using 41 genes that are targets of drugs identified in (A-C). The top 20 enriched GO terms are shown. (D) and (E) represent GO terms related to biological process and cellular component, respectively.

### Observing long-term drug responses of AML cells utilizing an optimized-microwell imaging system

We next attempted to observe the long-term effects of the predicted drugs affecting AML cell proliferation on cell cycle progression. From the top drugs affecting AML cell proliferation identified by DepMap data analysis (Effect size ≤-3.0) and clinically used drugs for AML treatment, we selected those involved in cell cycle and mitosis regulation. Firstly, from the 29 drugs, we selected 22 drugs whose targets were classified under the Top 3 Gene Ontology (GO) terms: cell cycle, cell cycle process, and mitotic cell cycle (Fig. 2A). Secondly, we broadly classified these 22 drugs into three groups based on their modes of action: Group 1 consisted of 8 drugs targeting DNA synthesis and chromatin organization, Group 2 comprised 11 drugs targeting mitotic progression, and Group 3 included 3 drugs targeting other processes. To examine the effects of drugs classified in Groups 1-3, we selected the TOP2A inhibitor idarubicin (IDR), the microtubule inhibitor vincristine (VCR) and the Nedd8-activating enzyme inhibitor pevonedistat (PVN), as a representative example of each group, respectively, because these drugs are clinically relevant for AML treatment^29–35^ (Fig. 2A).

**Figure 2.**
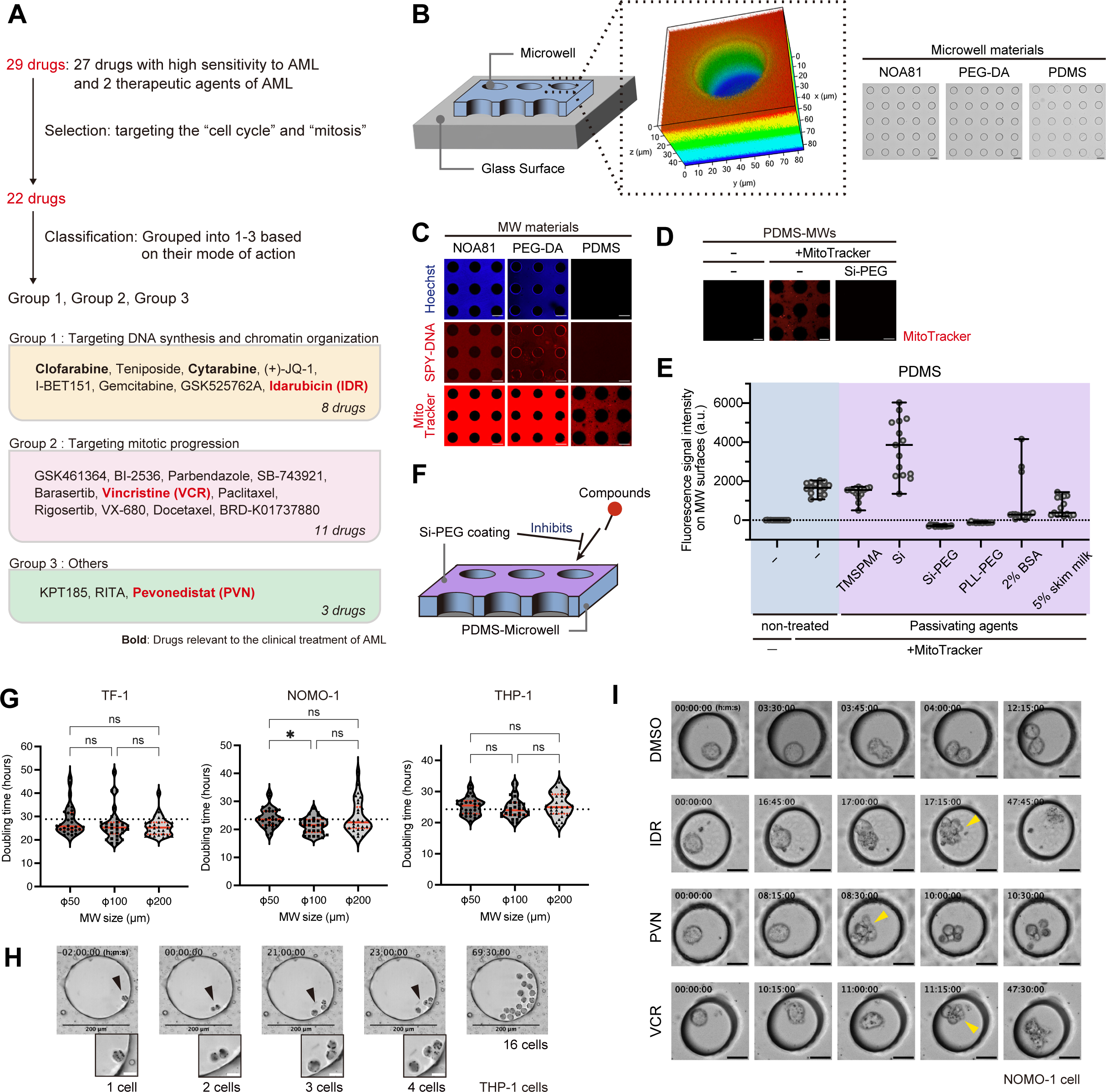
Optimized microwell system enables long-term observation of drug responses in AML cells. (**A**) Classification of the identified therapeutic agents based on their mechanisms of action. (**B**) The MWs were made on the glass surface of culture plates. A 3D-reconstituted image of a NOA81-based MW was shown in the middle panel. The autofluorescence of NOA81 was detected. Top-view images of microwell arrays fabricated with each material were shown on the right. (**C**) Adsorption of fluorescently-labeled chemicals on the MW resin surfaces. Scale bar: 100 μm. (**D**) Adsorption of MitoTracker on the PDMS MWs with or without silane-PEG passivation. Scale bar: 100 μm. (**E**) The average fluorescence signal intensities on the MW surfaces were quantified. n = 15 fields per each condition. Representative images were shown in Figure 2D and S2D. Data are shown as mean ± SD. (**F**) Schematic illustration of surface passivation of PDMS-MWs with Si-PEG to prevent the adsorption of compounds. (**G**) Violin plots showing the doubling time of individual AML cells in the Si-PEG-coated PDMS MWs. The black dashed line represents the doubling time in standard culture plates without MWs. The median is shown as a solid red line, and the first and third quartiles are shown as dashed red lines. n = 30 cells. Dunn’s multiple comparisons test was performed to obtain adjusted p-values. ns: not significant, *p < 0.05. (**H**) Time-lapse images of the proliferation of THP-1 cells in 200 μm diameter Si-PEG-coated PDMS MWs. Black and white scale bars: 200 μm and 20 μm, respectively. Arrowheads: magnified areas. (**I**) Time-lapse images of NOMO-1 cells treated with each drug in Si-PEG-coated PDMS MWs. Scale bar: 20 μm. Arrowheads: the timing of cell death.

To observe the effect of these therapeutic agents on AML cells, we attempted to perform long term live cell imaging. However, AML cells tend to flow in the medium due to suspension culture and easily drift out of the field of view during time-lapse imaging, making long-term observation challenging (THP-1 cells, Fig. S2A). To overcome this technical limitation, we applied microwells (MWs) for long-term live-cell observation in AML cells. To this end, we designed circular MWs with a height of 35 μm using several resins (NOA81^36^, PEG-DA^37^, and PDMS^38^, Fig. 2B). Furthermore, to properly evaluate the effects of therapeutic agents on cultured cells, it is important to prevent a decrease of chemical concentration in the culture medium due to adsorption on the MW surface^39,40^. To identify the optimal resin for MWs, we evaluated the adsorption of several fluorescently-labeled chemicals (e.g. MitoTracker) to the resins. The results showed that most compounds were hardly detected on the surface of the PDMS-based MWs, whereas they were strongly detected on the NOA81 and PEG-DA MWs (Fig. 2C, S2B-C). In addition, surface passivation of the PDMS MWs with Silane-PEG (Si-PEG) or PLL-PEG further suppressed adsorption of some chemicals on the surface (Fig. 2D-E, S2D-E). These results suggest that passivating the PDMS with Si-PEG or PLL-PEG could prevent a decrease of therapeutic agent concentration in the MW-based cell culture, enabling accurate analysis of therapeutic agent responses.

Using the Si-PEG coated PDMS MWs, we verified that the size of the MWs did not affect survival and doubling time of several AML cell lines (TF-1, NOMO-1 and THP-1, Fig. 2G, S2F) and enabled tracking of at least four cell divisions of individual cells over 72 hours (Fig. 2H). Furthermore, we successfully observed the accurate timing of cell death of AML cells induced by IDR, PVN and VCR (Fig. 2I). These results suggest that our optimized MW imaging system could be a powerful tool for studying the therapeutic agent response of AML cells.

### IDR, PVN, and VCR induce cell death from S/G2 to M phase in AML cells

To investigate the effects of IDR, PVN, and VCR (Fig. S3A-C) on cell cycle progression, we introduced the fluorescent, ubiquitination-based cell cycle indicator “Fucci”^41^ into AML cells, NOMO-1 and THP-1 cells. By using the Fucci-labelled AML cells and the optimized MW imaging system, we successfully observed multiple rounds of cell cycles in AML cells (Fig. 3A-B). In addition, the timing of the mitotic entry was judged by chromosome condensation by staining with SiR-DNA dye. Live-cell imaging showed that AML cells treated with IDR and PVN ultimately underwent cell death in either the S/G2 phase or mitosis (Fig. 3C-D), at approximately the IC_50_ concentration for each cell line (Fig. S3D-E). On the other hand, the treatment with VCR induced cell death in mitosis in most AML cells (Fig. 3C-D), at approximately the IC_50_ concentration for each cell line (Fig. S3F). These data indicate that these therapeutic agents induce cell death from S/G2 to M phase in AML cells. Importantly, we also found that IDR and PVN had an efficient cytotoxic effect on AML cells (MDS-L-2007, TF-1, NOMO-1 and THP-1 cells) in compared to non-hematopoietic cells (RPE-1, HeLa and U2OS cells), demonstrating that AML cells are particularly sensitive to these therapeutic agents (Fig. S3D-E). In addition, the percentage of aneuploid and polyploid cells with >4n DNA content increased with IDR and PVN treatment, which are not mitotic inhibitors like VCR, suggesting a disruption of mitotic processes by these therapeutic agents (Fig. S4A-D). In support of these findings, the IDR and PVN increased the percentage of cells with micronuclei in surviving cell populations (Fig. S4E-F). Furthermore, a trend toward increased nuclear volume was observed in the AML cells treated with these therapeutic agents (Fig. S4G). This phenotype can be explained by errors in the G2 phase or mitosis caused by IDR and PVN. Collectively, these data suggest that IDR, PVN and VCR disrupt cell cycle regulation, particularly during the S/G2 and M phase, and induce cell death in AML cells.

**Figure 3.**
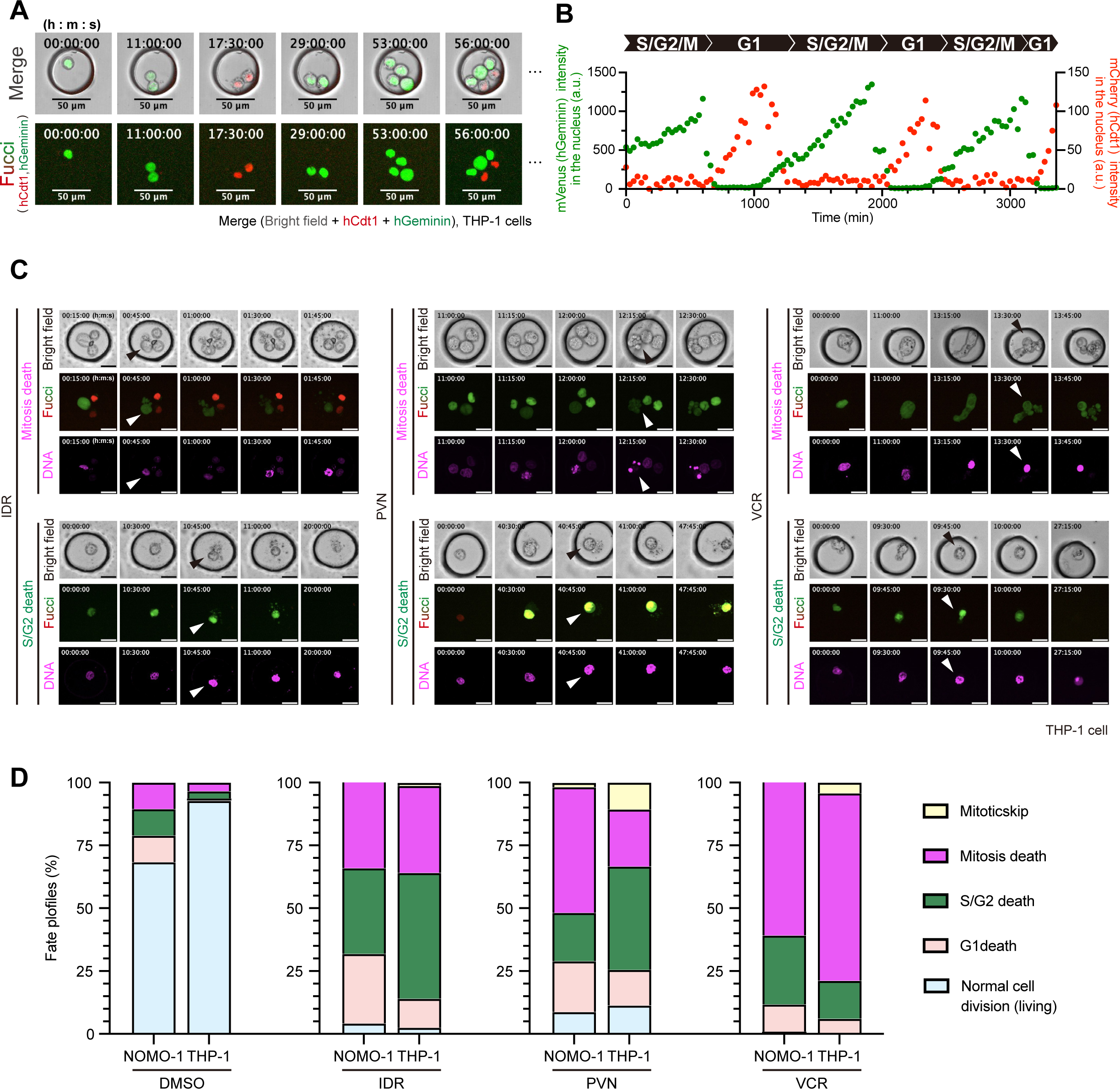
IDR, PVN, and VCR exert cytotoxic effects on AML cells during the S/G2 and M phases. (**A**) Time-lapse images of THP-1-Fucci cells stably expressing hGeminin-mVenus and hCdt1-mCherry in Si-PEG-coated PDMS MWs. Scale bars: 50 μm. (**B**) Temporal profiles of the mean fluorescence intensity of mCherry (red circles) and mVenus (green circles) shown in (A). (**C**) Time-lapse images showing cell death during S/G2 phase or mitosis, induced by 10 nM idarubicin (IDR), 40 nM pevonedistat (PVN), and 30 nM vincristine (VCR). Scale bar: 20 μm. Arrowheads: the timing of cell death. (**D**) Cell fate profiles for each cell line upon treatment with each drug at its IC_50_ concentration. n = 2 independent experiments, 114 (DMSO), 47 (IDR), 114 (PVN), and 102 (VCR) cells (NOMO-1 cell line), or 207 (DMSO), 78 (IDR), 141 (PVN), and 165 (VCR) cells (THP-1 cell line).

### AML cells are vulnerable to mitotic arrest

In actual therapeutic agent administration, not only the efficacy but also the rapid cellular response to a therapeutic agent at a constant concentration is a crucial factor, as the time to cell death is related to the cumulative exposure^42–44^. Therefore, we used the microwell-based observation system, which is an excellent method for quantitatively analyzing the speed of cellular response, to observe the time course of cell death induced by IDR and PVN treatment during the S/G2 phase and mitosis in AML cells, NOMO-1 and THP-1 cells. By measuring the time from cell cycle entry to cell death, we found that cell death in mitosis induced by IDR and PVN proceeded more rapidly than that in the S/G2 phase. Ninety percent of the mitotic cell death was induced within 1.5 hours on average (Fig. 4A). In line with this, it has been shown that non-hematopoietic cells tend to tolerate longer mitotic arrest^45,46^. Therefore, these results suggest that AML cells may be vulnerable to mitotic arrest as compared to non-hematopoietic cells. To check this possibility, we compared the time course of mitotic cell death induced by vincristine among AML and non-hematopoietic cells. In non-hematopoietic cell lines (RPE-1 and HeLa), cell death occurred during a prolonged mitotic arrest and progressed gradually over 20 hours until all cells died (Fig. 4B-D). In contrast, in AML cells, all mitotic cell death occurred rapidly within 5 hours (1.5 hours on average in Fig. 4B-D). These data strongly suggest that AML cells are more vulnerable to mitotic arrest than non-hematopoietic cells.

**Figure 4.**
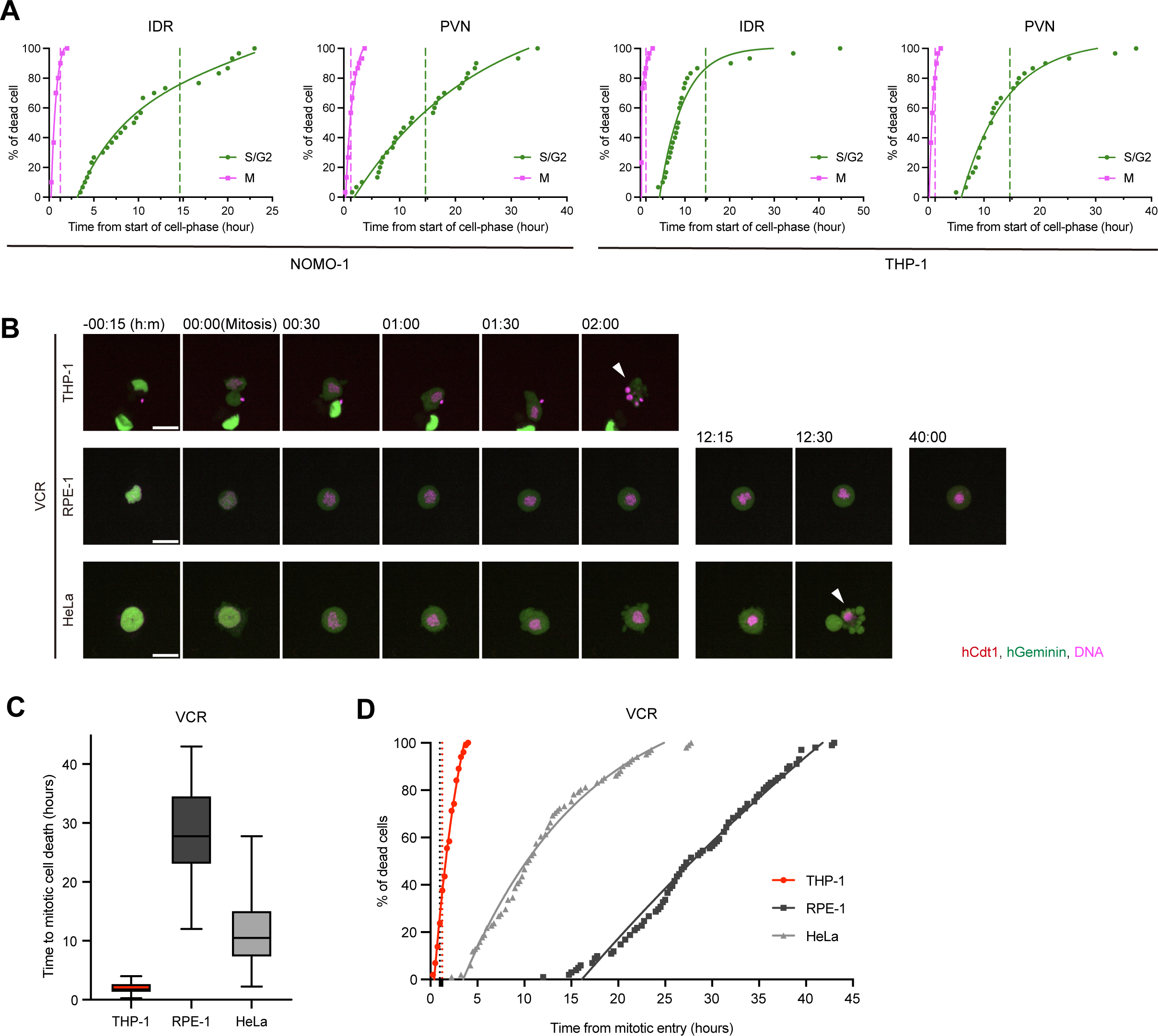
AML cells have low tolerance to mitotic arrest. (**A**) The time required for cell death from the onset of a particular cell-cycle phase. Each plot shows the cumulative percentage of cell death at each time point. n = 30 cells. (**B**) Time-lapse images showing cell death of cells arrested in mitosis by 30 nM vincristine (VCR). Arrowheads: the timing of cell death. (**C**) Box-whisker plot showing the time required for cell death from the mitotic onset in the presence of VCR (30 nM). The maximum observation time is 48 hours. n > 100 cells from 2 independent experiments. (**D**) The time required for cell death from the mitotic onset. Each plot shows the cumulative percentage of cell death at each time point in (C). n > 100 from 2 independent experiments.

### Forced mitotic entry of IDA or PVN-treated AML cells enhances the cytotoxicity

Our time-lapse imaging analysis revealed that the mitotic cell death induced by IDR or PVN proceeded more rapidly than that in the S/G2 phase (Fig. 4A) and the IDR- or PVN-treated cells exhibited a prolonged S/G2 phase. Therefore, we expected that releasing the IDR- or PVN-treated AML cells from S/G2 arrest and promoting their mitotic entry would induce efficient cell death during mitosis. It has been suggested that the G2/M checkpoint, which is regulated by the phosphorylation of Chk1 by ATR and ATM, arrests cells in S/G2 phase^47^. We found that the treatment with IDR and PVN increased the levels of the phosphorylation of Chk1 (Fig. 5A), as well as the levels of CDK1 and Cyclin B1 proteins, which are known to be upregulated in the G2 phase (Fig. 5A). These results suggest that IDA and PVN induce cell cycle arrest by activating the G2/M checkpoint in AML cells. Hence, we examined the effects of Chk1 inhibitor CHIR124^48,49^ to release the cells from S/G2 arrest and promote mitotic entry (Fig. 5B). Live-cell imaging showed that cell cycle arrest at the S/G2 was alleviated in THP-1 cells upon treatment with CHIR124 at non-cytotoxic concentration (Figure S5A), and that the THP-1 cells released from the S/G2 arrest underwent cell death during mitosis (Fig. 5C). Furthermore, CHIR124 was effective in inducing apoptosis in combinations with IDR and PVN (Figure 5D, S5B-D). These results demonstrate that the release of cells from the G2/M checkpoint by CHIR124 promotes their mitotic entry and enhances the cytotoxicity of IDR and PVN in AML cells.

**Figure 5.**
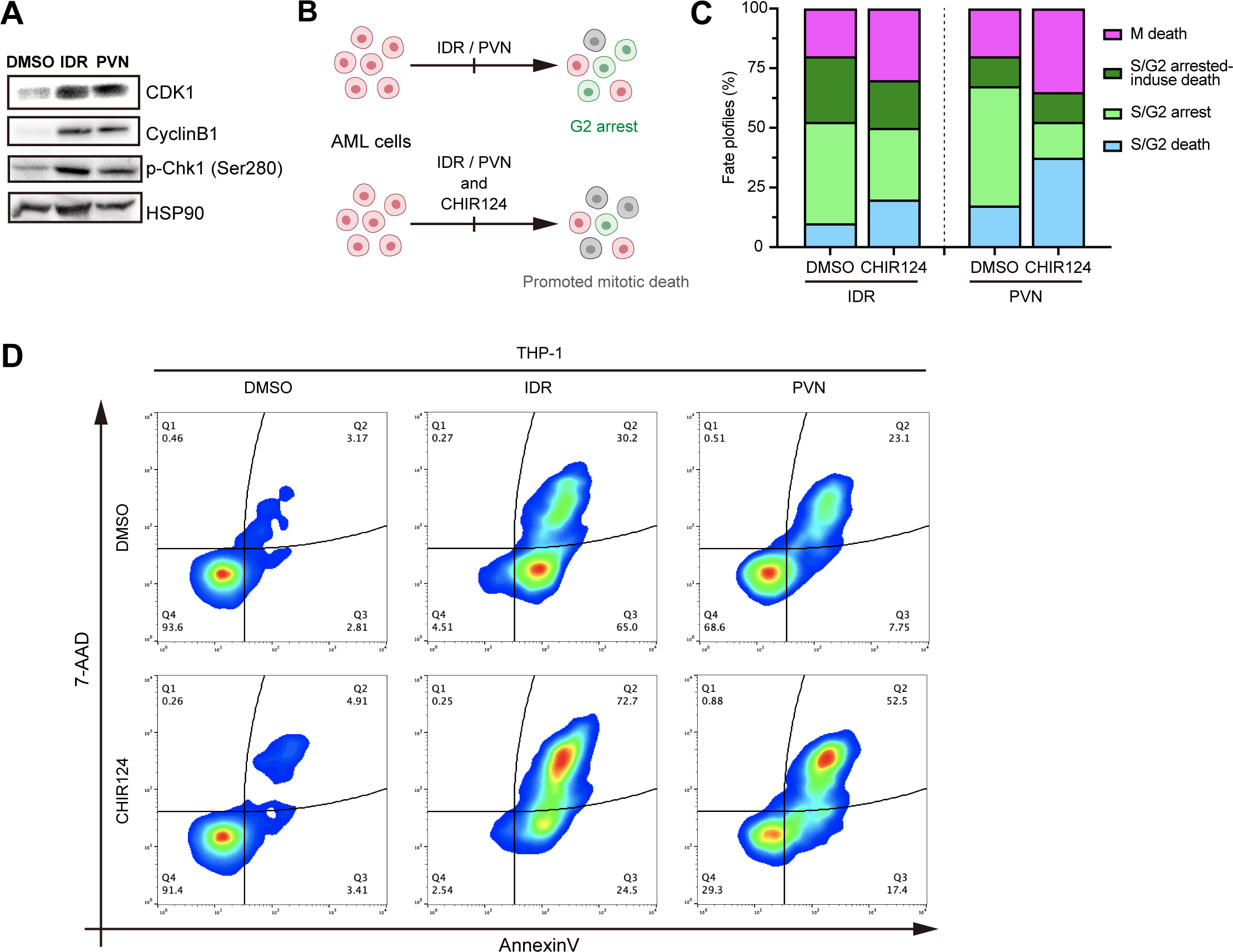
Combined treatment of IDR or PVN with the Chk1 inhibitor CHIR124 effectively induces cell death. (**A**) Expression levels of CDK1, CyclinB1, and p-Chk1 (Ser280) proteins in NOMO-1 cells treated with 10 nM idarubicin (IDR) and 40 nM pevonedistat (PVN) for 48 hours. (**B**) Schematic of the forced mitotic entry in IDR- or PVN-treated AML cells by suppressing the G2/M checkpoint with a Chk1 inhibitor. (**C**) Cell fate profiles for THP-1 cells upon treatment with 3 nM idarubicin (IDR), 30 nM pevonedistat (PVN), and 3 nM CHIR124. n = 40 cells. Based on (Fig 4A), cells that were stalled in the S/G2 phase for 16 hours were defined as S/G2 arrest, and a distinction was made between those that subsequently underwent cell death and those that remained arrested until the end of the observation period. (**D**) Heatmap of apoptosis status measured by flow cytometry in THP-1 cells treated with IDR and PVN alone and in combination with CHIR124. CHIR124 was used at a concentration that does not affect cell toxicity over 48 hours. More than 5000 cells were detected. The heatmap was divided into four quadrants (Q1-Q4), and the cells in each quadrant were displayed as percentiles up to 100. Q1 (non-apoptotic cell death: − AnnexinV/+7-AAD), Q2 (late apoptosis/Dead: +AnnexinV/+7-AAD), Q3 (early apoptosis: +AnnexinV/−7-AAD), and Q4 (living: −AnnexinV/−7-AAD).

## Discussion

Induction therapy for AML is currently based on chemotherapy, and its efficacy has a significant impact on therapeutic outcome^42–44^. However, AML is recognized as a disease with poor treatment outcomes, and providing new AML treatment strategies to improve these outcomes is an urgent challenge. Therefore, understanding the vulnerabilities of AML cells is crucial for developing effective treatment strategies for AML. In this study, we elucidated the vulnerability of AML cells induced by therapeutic agents. The informatics analysis using DepMap data predicted that AML cells tend to be highly sensitive to drugs that target cell cycle progression and mitosis (Fig. 1D). Several studies using genome-wide CRISPR-Cas9 knockout screens have been conducted to investigate vulnerabilities in cancer. However, these approaches cannot evaluate core essential genes for cell proliferation. Our approach of comparing drug sensitivity provides a new avenue to overcome these limitations and identify vulnerabilities in AML that directly affect cell proliferation events, such as the cell cycle and mitosis. The MW-based imaging system, optimized for drug analysis, revealed that drugs targeting these processes, such as IDR, PVN, and VCR, efficiently and rapidly induced cell death during mitosis in AML cells (Fig. 3D, 4A,D). These results highlight a latent vulnerability in the processes of mitosis in AML cells, suggesting that targeting this vulnerability could be an effective therapeutic strategy. Further studies based on these findings will lead to the development of new therapeutic agents targeting AML.

IDR, a widely used therapeutic agents in current AML treatment, is known to exert its antitumor effects by intercalating into DNA and inhibiting topoisomerase II^50^. Therefore, it is reasonable to expect that IDR induces cell death in the processes of DNA replication during the S-G2 phase. However, our study revealed that IDR also strongly induces mitotic catastrophe in AML cells (Fig. 3C-D). This suggests that DNA damage caused by IDR may induce vulnerability in AML cells during mitosis. Further analysis revealed that cell death induced by IDR was triggered more efficiently and rapidly in mitosis compared to the S/G2 phase (Fig 4A). These results indicate that the mitotic system in AML cells is vulnerable to the defects in DNA replication processes, and targeting these processes may be a promising strategy for AML treatment.

PVN has been reported to be effective against TP53-mutated AML and secondary AML (s-AML), which are known to have a poor prognosis and are refractory to treatment, based on the results of a phase II clinical trial^31^. The MDS-L-2007^51^ and TF-1^52^ cells used in this study are both from MDS or s-AML. Notably, high-risk s-AML and MDS are frequently characterized by a complex karyotype, which arises from incomplete cell division. The presence of a complex karyotype is a significant factor in classifying MDS as high-risk according to the current clinical prognostic score (IPSS-R)^53^. This suggests that high-risk s-AML and MDS may have inherent vulnerabilities in their mitotic processes, making them potentially more susceptible to therapeutic agents targeting mitotic processes. This notion is further supported by a recent report demonstrating that decitabine, a DNA methylation inhibitor, induces cytotoxicity in s-AML cells at clinically relevant concentrations by disrupting chromosome segregation^20^. Although decitabine and PVN have different modes of action, both therapeutic agents affect mitosis, leading to similar phenotypic outcomes. Therefore, by elucidating the mechanisms of chromosome segregation during mitosis in s-AML cells, it may be possible to exploit these vulnerabilities and develop effective therapies for s-AML, which currently has a poor prognosis.

In cancer treatment, the dosage and duration of chemotherapy drug administration are limited by acute and cumulative toxicities in normal tissues^42–44^. Moreover, the use of therapeutic agents at ineffective concentrations can cause epigenetic changes in tumor cells, leading to the acquisition of disease-related drug resistance^54–56^. Our study has demonstrated that the combination of a Chk1 inhibitor with therapeutic agents such as IDR and PVN at lower concentrations can induce early cell death during the M phase in AML cells. These cell cycle-targeted approaches are likely to be more effective against cells with shorter doubling times. Moreover, it is known that blood cells account for the majority of daily cellular turnover in the human body^57^, suggesting that these approaches may be particularly effective against hematological malignancies, which often have even shorter doubling times in their malignant state. This targeted approach may potentially reduce the required dosage and duration of therapeutic agent administration, which could minimize side effects on normal tissues. Furthermore, by inducing rapid cell death in AML cells, this strategy could effectively eliminate tumor cells before they have acquired therapeutic resistance. Therefore, further evaluation of these approaches, which may lead to more effective and safer AML treatments, would be valuable for improving therapeutic strategies for AML.

## Acknowledgements

We thank the Broad Institute for providing the DepMap dataset and all members of Kitagawa laboratory for fruitful discussion. This work was supported by the Japan Society for the Promotion of Science (19H05651 [D.K.], 20K22701 [S.H.], 21H02623 [S.H.], 22K19305 [M.F.], 22K19370 [S.H.], 22K20624[S.Y.], 23H02627 [T.C.], 23K14176[S.Y.], 24K02174 [S.H.]), Japan Science and Technology Agency (JPMJCR22E1[D.K.], JPMJPR21EC [S.H.]), Takeda Science Foundation [D.K., T.C., S.Y.], Princess Takamatsu Cancer Research Fund [D.K., S.Y.], Uehara Memorial Foundation [T.C., S.Y.], Inamori Foundation [S.Y.], Naito Foundation [T.C.], Kanae Foundation for the Promotion of Medical Science [T.C.], Kato Memorial Bioscience Foundation [S.H.], Mochida Memorial Foundation for Medical and Pharmaceutical Research [T.C., S.H.], Nakajima Foundation [T.C.], and Sumitomo Foundation [T.C.], Nippon Shinyaku Research Grant program [T.C.], Astellas Foundation for research on metabolic disorders [S.Y.], Kishimoto Fund Research Grant from the Senri Life Science Foundation [S.Y.] and Noguchi Shitagau Research Grant from the Noguchi Institute [S.Y.].

## Author contributions

Conceptualization: R.N., S.Y., T.C. and D.K.; Methodology: R.N., T.Y., S.Y., H.S.; Reagents and Cell lines; T.Y., S.G., T.K.; Formal analysis: R.N.; Investigation: R.N.; Data curation: R.N.; Visualization: R.N.; Writing (original draft), R.N.; Writing (modification): S.Y., T.C. and D.K.; Supervision: S.Y., T.C. and D.K. Project administration: R.N., S.Y., T.C. and D.K.; Funding acquisition: S.Y., T.C., M.F., S.H. and D.K. All authors contributed to discussions and manuscript preparation.

## Materials and Methods

### Informatics analysis

A comparative analysis of drug sensitivity between AML cells and non-HM cells was performed using the Drug sensitivity AUC (CTD^2^) data registered in the DepMap database. In addition, drugs that significantly altered sensitivity in AML and non-HM cells with an Effect size indicating drug sensitivity of ≤-3.0 (AML sensitive) or ≥0 (non-HM sensitive) and a P value of ≤0.01 were selected. The targets of each drug were extracted by referring to the Drug Repurposing Hub (https://repo-hub.broadinstitute.org/repurposing-app) and Cancer Therapeutics Response Portal v2 (https://portals.broadinstitute.org/ctrp.v2.1/). For the targets, the common targets registered in both databases were extracted, and if a target was registered in only one database, these targets were also extracted. For the targets of docetaxel, nine isotypes of TUBB (TUBB, TUBB1, TUBB2A, TUBB2B, TUBB3, TUBB4A, TUBB4B, TUBB6, TUBB8) were extracted as target molecules and therefore combined as TUBB. Furthermore, data mining was performed by conducting an unbiased Gene Ontology (GO) enrichment analysis using g.Profiler (https://biit.cs.ut.ee/gprofiler/gost) on the extracted targets, which yielded the Biological Process (BP) and Cellular Component (CC) categories that these targets are involved in.

### Cell culture

NOMO-1 (JCRB Cell Bank, IFO50474) and THP-1 (ATCC, TIB-202) cells were cultured in RPMI-1640 (Nacalai Tesque, 30264-56) medium supplemented with 10% fetal bovine serum (FBS; NICHIREI, 175012) and 100 U/mL penicillin and 100 μg/mL streptomycin (1% P/S; Nacalai Tesque, 09367-34). MDS-L-2007 cells^51^ were cultured in RPMI-1640 medium supplemented with 10% FBS, 1% P/S and 2 ng/mL human IL-3 (PeproTech, AF-300-23). TF-1 cells^52^ were cultured in RPMI-1640 medium supplemented with 10% FBS, 1% P/S and 2 ng/mL human G-CSF (PeproTech, AF-200-03). RPE-1 (ATCC, CRL-4000), HeLa (ECACC, 93021013), U2OS (ECACC, 92022711) and HEK293T (ECACC, 12022001) cells were authenticated by the suppliers via short tandem repeat profiling before purchase. RPE-1 cells were cultured in DMEM/F-12 medium (Nacalai Tesque, 11581-15) supplemented with 10% FBS and 1% P/S. HeLa and HEK293T cells were cultured in Dulbecco’s modified Eagle’s medium (DMEM; Nacalai Tesque, 08459-64) supplemented with 10% FBS and 1% P/S. U2OS cells were cultured in McCoy’s 5A (Modified) Medium (Nacalai Tesque, 16600082) supplemented with 10% FBS and 1% P/S. All cells were maintained at 37°C in a 5% CO_2_ atmosphere.

### Generation of cell lines by retroviral-mediated integration

FUCCI-expressing AML cell lines were established by retroviral-mediated integration. Following plasmids were mixed with 250 mM CaCl_2_ in MilliQ: 10 μg of packing plasmid (M57), 3 μg of envelope plasmid (RD114), and 12 μg of transfer plasmids (pMXs mVenus-hGeminin (1/110) or pMXs-IB mCherry-hCdt1 (30/120)). The mixture was then diluted in filtered HeBS (0.28 M NaCl, 49.7mM HEPES, 1.5 mM Na_2_HPO_4_·12H_2_O, pH: 7.2) and transfected into HEK293T cells using the calcium phosphate method. The medium was harvested and filtered through a 0.45 μm filter (Merck Millipore, SLHVR33RS). AML cells were infected with the virus-containing medium in the presence of Retronectin (Takara, #T100A), according to the manufacturer’s protocol. Single-cell clones were isolated by using a FACS AriaIII/IIIu (BD).

### Chemical compounds

The following chemical compounds were used in this study: Idarubicin (IDR; Selleck, #S1228), Pevonedistat (PVN; MedChemExpress, #HY-10484), Vincristine (Cayman chemical, #11764), VE821 (Selleck, #S8007), CHIR124 (Cayman chemical, #16553). All compounds were dissolved in DMSO according to the manufacturer’s guidelines and stored at -20°C (or -80°C for long-term storage).

### Cell viability assay

AML cell lines (2 × 10^4^ or 3 × 10^4^ cells/well in 96-well plates) and non-hematopoietic cell lines (5 × 10^3^ cells/well in 96-well plates) were treated with each agent (final DMSO concentration was 1%) for two days. 5 or 10 μL of Cell Count Reagent SF (Nacalai Tesque, 07553) was added to the culture. After 2–5 hours incubation, the absorbance at 450 nm was measured with the FLUOstar OPTIMA-5 microplate reader (BMG LabTech), and the cell viability (control %) was determined. IC_50_ values were calculated from fitting curves of cell viability using the GraphPad Prism 9 software.

### Cell cycle analysis

The cells were washed once with PBS and fixed with 70% EtOH at -30°C for 5–15 hours. After fixation, the cells were washed with PBS and thoroughly mixed with the Muse™ Cell Cycle Kit (Cytek Biosciences, # MCH100106). The mixture was then incubated at room temperature for at least 30 minutes. Subsequently, the sample was diluted with PBS and analyzed using the Guava^®^ Muse™ Cell Analyzer. Analyses were performed with FlowJo Software (Tree Star Inc.)

### Nuclear volume/morphology analysis

The cells were fixed in 4% paraformaldehyde in phosphate-buffered saline (PFA) (Nacalai Tesque, #09154-85) for 15 minutes at room temperature and washed several times with PBS. The cells were then resuspended in PBS, and nuclei were stained by adding 0.4 μg/ml Hoechst 33258 (Nacalai Tesque, #19173-41). The cell suspension was added to a 35 mm glass-bottom dish (Greiner Bio-One, #627871). Z-stack images were acquired using a Leica TCS SP8 confocal microscope (Leica), ensuring that all nuclei were captured with a z-step of 0.15 μm.

### Western Blotting

Cells were lysed on ice in RIPA buffer (150 mM NaCl, 15 mM MgCl2, 5 mM EDTA, 1% (v/v) NP-40, 0.5%(w/v) Sodium Deoxycholate, 0.1% (v/v) SDS, 50 mM Tris-HCl (pH 8.0), 0.1% (v/v) Protease Inhibitor Cocktail (Nacalai Tesque, #25955-11), and 0.1% (v/v) Phosphatase Inhibitor Cocktail (Nacalai Tesque, #07575-51)). The total protein concentration of cell lysate was then equalized using the Protein Assay BCA Kit (Nacalai Tesque #06385-00). Protein samples in SDS sample buffer (Nacalai Tesque, 09499-14) were subjected to SDS-PAGE on a 10% polyacrylamide gel and subsequently transferred to Immobilon-P PVDF-membrane (Merck Millipore, IPVH85R). The membrane was blocked with 5% (w/v) skimmed milk in PBS containing 0.02% (v/v) Tween (PBS-T) for 30 min at room temperature or Blocking One P (Nacalai Tesque, #05999-84) for 20 min and then washed three times with PBS-T. The membrane was incubated with primary antibodies in 5% BSA and 0.01% NaN_3_ in PBS-T overnight at 4°C. After washed three times with PBS-T, the membrane was incubated with secondary antibodies in 5% skimmed milk in PBS-T for 1 hour at room temperature. The membrane was washed three times with PBS-T before chemiluminescent detection. Signal detection was carried out with Chemi-Lumi One L (Nacalai Tesque, 07880), Chemi-Lumi One Super (Nacalai Tesque, 02230), or Chemi-Lumi One Ultra (Nacalai Tesque, 11644) using a ChemiDoc XRS Plus (Bio-Rad).

### Construction of microwells

The master mold (approximately 35 μm in thickness) was fabricated through photolithography using SU8-3025 (Kayaku Advanced Materials, #311072), which was exposed to UV light (Polarstar, 30 W, 6 sec) and then vapor silanized with Trichloro (1H,1H,2H,2H-perfluoro-octyl) silane (Si; Sigma, 448931). To make the 1st PDMS, a mixture of prepolymer and curing agent (Dow, SYLGARD 184 silicone elastomer kit) was poured onto the master mold. It was baked at 95 °C for 1.5 hours. The 1st PDMS was then vapor silanized with Trichloro (1H,1H,2H,2H-perfluoro-octyl) silane. The 2nd PDMS was fablicated from the silanized 1st PDMS as a template. The 2nd PDMS was cut into small pieces and used as PDMS stamps. To fabricate microwells on the culture surface, the PDMS stamp was placed with the pillar surface facing the glass substrate. The microwell resin (NOA81, PEG-DA or PDMS) was dispensed onto the side of the PDMS stamp, allowing the resin to fill the spaces between the PDMS pillars by capillary action, and then cured as follows. NOA81 was cured by exposure to UV light for 15 sec. In the case of PEG-DA fabrication, Poly(ethylene glycol)diacrylate (Sigma, #475629) was mixed with 2-Hydroxy-2-methylpropiophenone (Sigma, #405655) at 5%(v/v) as a photopolymerization initiator. The PEG-DA mixture was cured by exposure to UV light for 45 sec. When fabricating MWs with PEG-DA, the culture chambers coated with 3-(Trimethoxysilyl) propyl methacrylate (TMSPMA; Sigma, 440159) were used. To prepare PDMS-based microwells, the mixture of prepolymer and curing agent was cured by heating at 95 °C for 1 hour. For live cell imaging, the PDMS-based microwells were subjected to surface passivation. The microwell-containing culture plates were incubated with Si-PEG solution (1 mg/ml of mPEG5K-Silane (Sigma-Aldrich, JKA3037-1G) in 99% ethanol and 0.1% HCl) at room temperature overnight to prevent non-specific binding of chemical compounds to the microwell surface.

### Quantification of adsorption of compounds on MW

The non-specific adsorption of compounds on MW was assessed using fluorescent probes. The following fluorescent probes were used in this study: SPY555-DNA (Cytoskeleton, Inc, #CY-SC201), SPY555-tubulin (Cytoskeleton, Inc, #CY-SC203), SiR-DNA (2 nM, Cytoskeleton, Inc, #CY-SC007), SiR-Tubulin (20 nM, Cytoskeleton, Inc, #CY-SC002), SiR-Actin (5 nM, Cytoskeleton, Inc, #CY-SC001), Hoechst33258 (1 μg/mL), PlasMem Bright Green (10 μg/mL, FUJIFILM Wako, #P504), MitoTracker (1 μM, Invitrogen, #M7512), Mito-Bright LT Green (1 μg/mL, DOJINDO, #MT10), LysoTracker (1 μM, Invitrogen, #L7526), or GOLGI ID^®^ Green (1 μg/mL, Enzo Life science, #ENZ-51028-K100). SPY555-DNA and SPY555-tubulin were used at a 5-fold dilution from the manufacturer’s protocol, which corresponds to the actual staining concentration. MWs with a diameter of 100 μm were fabricated on a 24-well glass-bottom culture plate (Greiner Bio-One, #662892). Then, 500 μL RPMI-1640 medium and fluorescent probes were added and the plate was incubated at 37°C with 5% CO2 for 2-3 hours. Z-stack images (Range: 60 μm, Slice: 30 frames, Step: 2 μm) including the top and bottom surfaces of the MW were acquired using a confocal quantitative image cytometer CellVoyager CQ1 (YOKOGAWA, Japan). Each excitation wavelength was as follows: 405 nm for Hoechst 33258, 488 nm for PlasMem Bright Green, Mito-Bright LT Green, LysoTracker, or GOLGI ID® Green, 561 nm for SPY555-DNA, SPY555-tubulin, or MitoTracker, and 640 nm for SiR-DNA, SiR-Tubulin, or SiR-Actin. The maximum intensity projection images were analyzed using Fiji (NIH) by creating a circle with a diameter of 110 μm to include the MW. The mean pixel intensities within the circular region were measured, and the intensities of the glass bottom surfaces outside the region where the MWs exist were subtracted as background for quantification.

### Surface passivation of microwells

The following agents were evaluated for surface passivation of microwells: TMSPMA, Si, Si-PEG, PLL-PEG (0.1 mg/mL in 10 mM HEPES buffer, Nanosof, #11354-x=150-2000-30%), BSA (Sigma-Aldrich, #A9647), or skim milk (5% (w/v) in PBS-T). For treatment with Si and TMSPMA, the microwells were incubated with vaporized gas of TMSPMA or Si under vacuum conditions for a minimum of 3.5 hours. For Si-PEG treatment, the microwells were incubated with the Si-PEG in EtOH containing hydrochloric acid under light-shielded conditions overnight. In the case of PLL-PEG, the microwells were incubated with 10 mM PLL-PEG in HEPES (KOH) buffer at pH 7.5 for 1 hour. Passivation with BSA was performed by incubating the microwells with 2% BSA in PBS for at least 1 hour. Skim milk treatment was performed by incubating the microwells with 5% skim milk in PBS-T for at least 1 hour.

### Live cell imaging

For live cell imaging using a Confocal Scanner Box CellVoyager CQ1 (YOKOGAWA, Japan), a custom-made PDMS-MW culture chamber on a 24-well glass-bottom plate (IWAKI, #5826-024 or Greiner-bio-one, #662892) was used. The chamber was passivated with Si-PEG O/N at room temperature. The stage incubator was maintained at 37°C with 5% CO2, and Z-stack images (Range: 18.0 μm, Slice: 14 images, Step: 1.4 μm, Binning: 2) were acquired every 15 minutes for over 48 hours using a 40× dry objective lens. For live cell imaging using a Spinning disk confocal scanner box CellVoyager CV1000 (YOKOGAWA, Japan), a custom-made PDMS-MW culture chamber on a 35-mm glass-bottom dish (Greiner Bio-One, #627871) was used. The chamber was passivated with Si-PEG at room temperature overnight. The stage incubator was maintained at 37°C with 5% CO2, and Z-stack images (Range: 17.0 μm, Slice: 14 images, Step: 1.3 μm, Binning: 2) were acquired every 15 minutes for over 48 hours using a 60× oil-immersion objective lens. Maximum intensity projections of representative images were created using Fiji. For cell cycle progression analysis, AML cells transduced with pMXs mVenus-hGeminin (1/110) (G1 phase) and pMXs-IB mCherry-hCdt1 (30/120) (S/G2/M phase) were stained with SiR-DNA (20 nM) and cultured in the MW. The timing of cell death was determined based on cell rupture or morphological abnormalities in bright-field images and a sudden increase in the fluorescence intensity of the SiR-DNA signal.

### Apoptosis analysis

The cells were washed with PBS and thoroughly mixed with the Muse™ Annexin V & Dead Cell Kit (Cytek Biosciences, #MCH10015) in PBS. The cells were then incubated at room temperature for at least 20 minutes. Subsequently, the cells were diluted with pure water and were analyzed using the Guava^®^ Muse™ Cell Analyzer. Analyses were performed with FlowJo Software (Tree Star Inc.)

---

**Figure S1.**
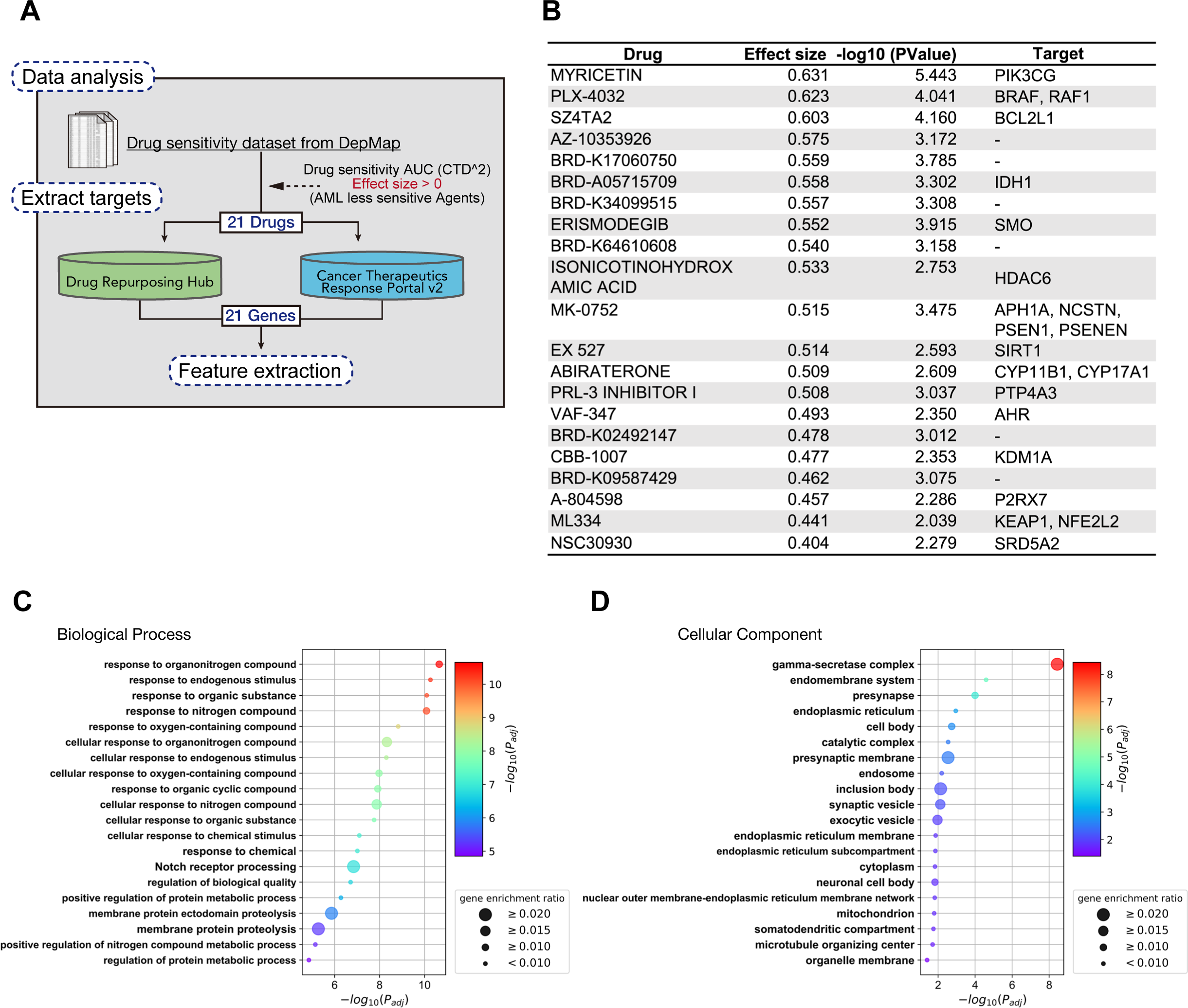
DepMap data analysis predicted drugs targeting organonitrogen compound response and gamma-secretase complex to be less effective against AML cells. **(A)** Identification of less effective drugs for AML cells through comparison of drug sensitivity AUC (CTD^2) data from DepMap. (**B**) Table of drugs classified as less sensitive to AML cells based on (Figure 1B) and their targets. (**C-D**) Gene Ontology (GO) enrichment analysis using g.Profiler was conducted using 21 genes that are targets of drugs exhibiting a trend of reduced sensitivity to AML cells. The top 20 enriched GO terms are shown. (C) and (D) represent the GO terms related to biological process and cellular component, respectively.

**Figure S2.**
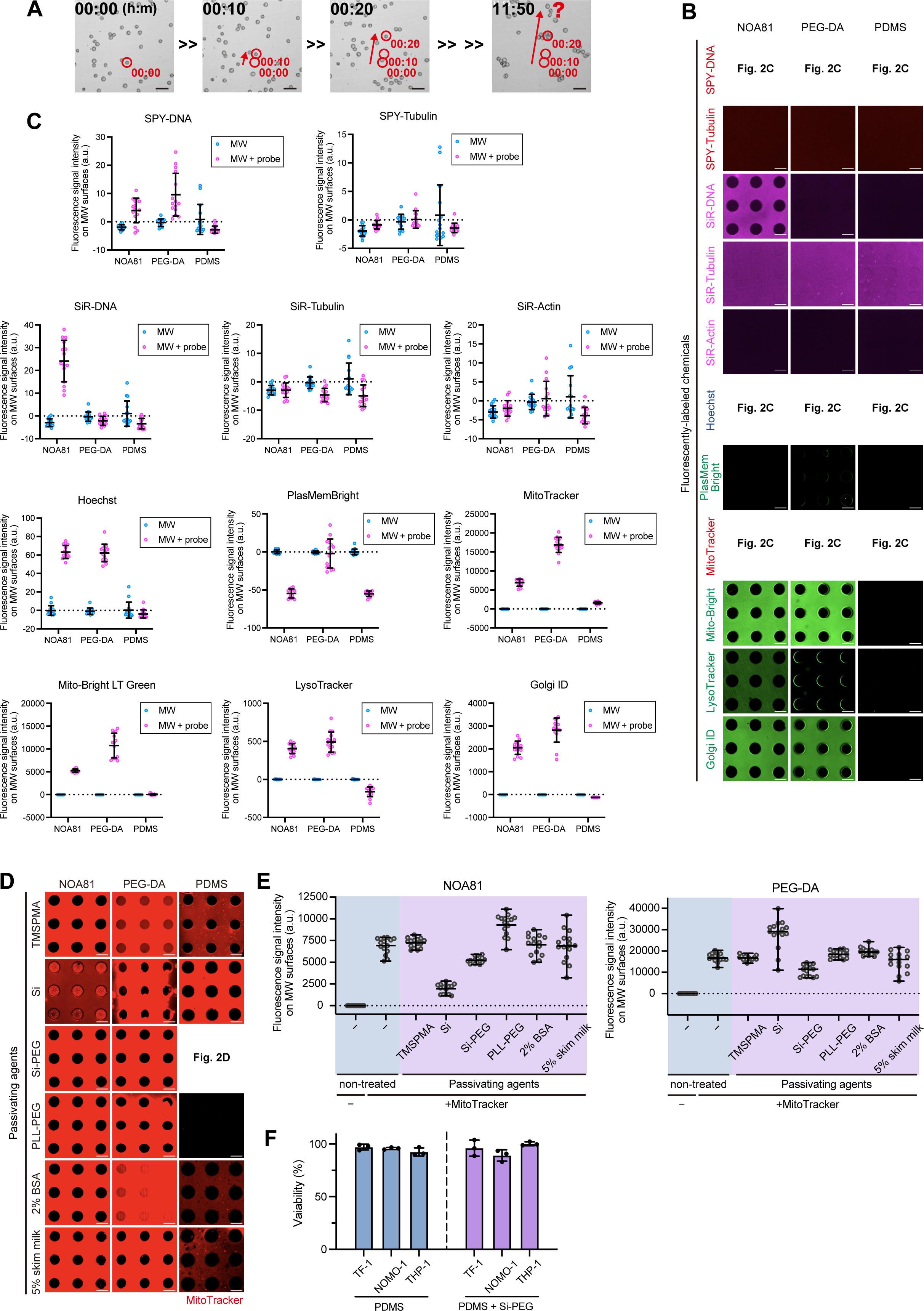
Optimization of a MW-based imaging system for monitoring drug responses in AML cells. (A) Time-lapse images of THP-1 cells drifting out of the field of view due to flow under standard culture conditions. Scale bar: 20 μm. Red arrows indicate the movement of cells from the initial time point. (**B**) Representative images of different fluorescently-labeled chemicals adsorbed on the MW resin surface. Scale bar: 100 μm. (**C**) The average fluorescence signal intensities were quantified within a 110 μm circular region containing MW from 15 randomly selected fields. Data are shown as mean ± SD. (**D**) Representative images of the MitoTracker adsorption on each passivated MW resin surface. Scale bar: 100 μm. (**E**) The average fluorescence signal intensities were quantified within a 110 μm circular region containing MW from 15 randomly selected fields. Data are shown as mean ± SD. (**F**) Cell viability of each cell type in the presence of PDMS or Si-PEG coated PDMS. Cell survival was determined by a cell proliferation assay after 48 hours of culture. The cell viability in an untreated condition was set as 100% viability.

**Figure S3.**
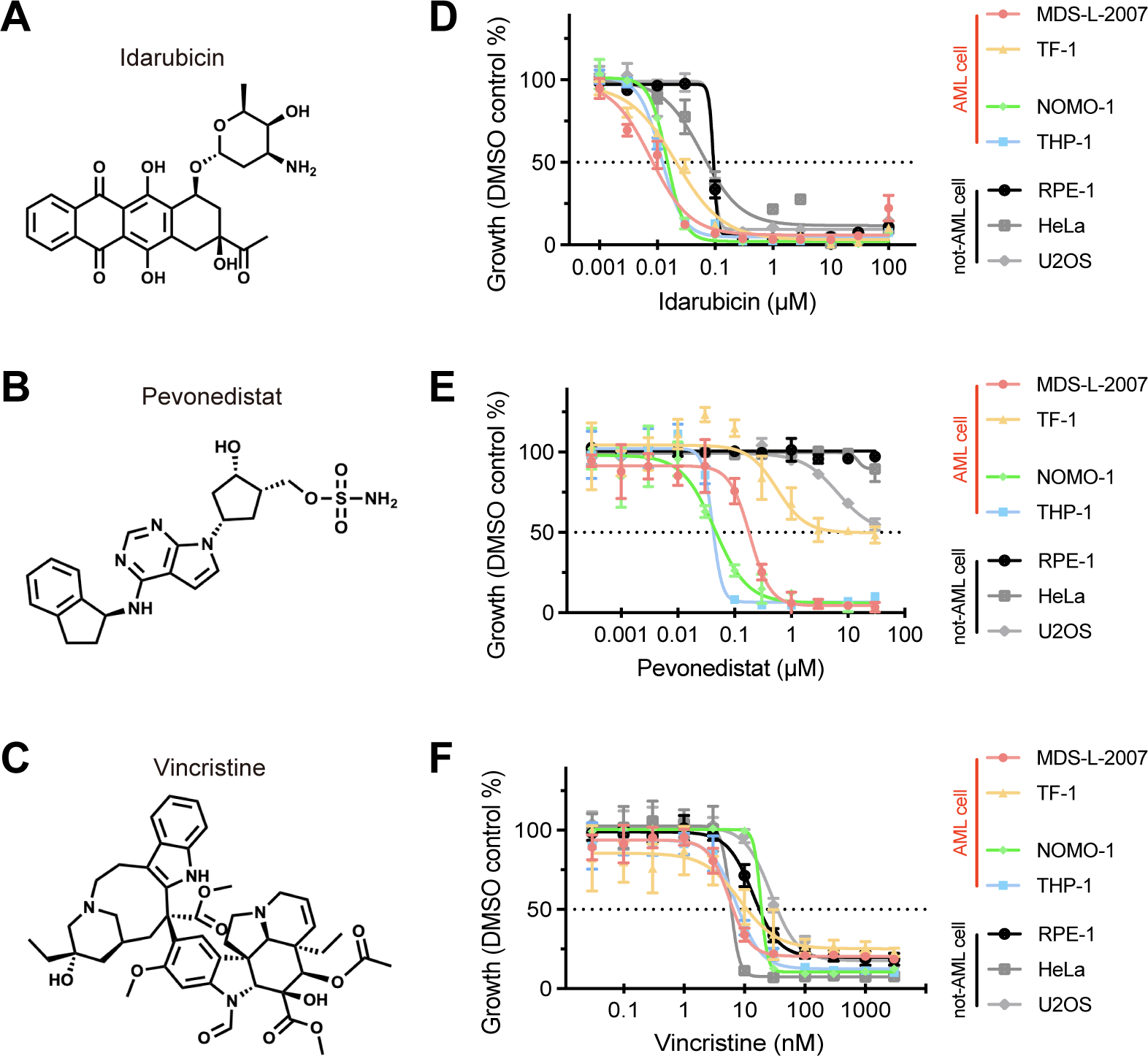
Validation of the cytotoxic concentrations of IDR, PVN, and VCR for each cell line. **(A)** Chemical structure of idarubicin (IDR). **(B)** Chemical structure of pevonedistat (PVN). (**C**) Chemical structure of vincristine (VCR). (**D**) Cell survival curves in response to IDR for each cell line. (**E**) Cell survival curves in response to PVN for each cell line. (**F**) Cell survival curves in response to VCR for each cell line. (**D-F**) The cell survival curves were determined by a cell proliferation assay after 48 hours of drug treatment. n>3. Data are shown as mean ± SD.

**Figure S4.**
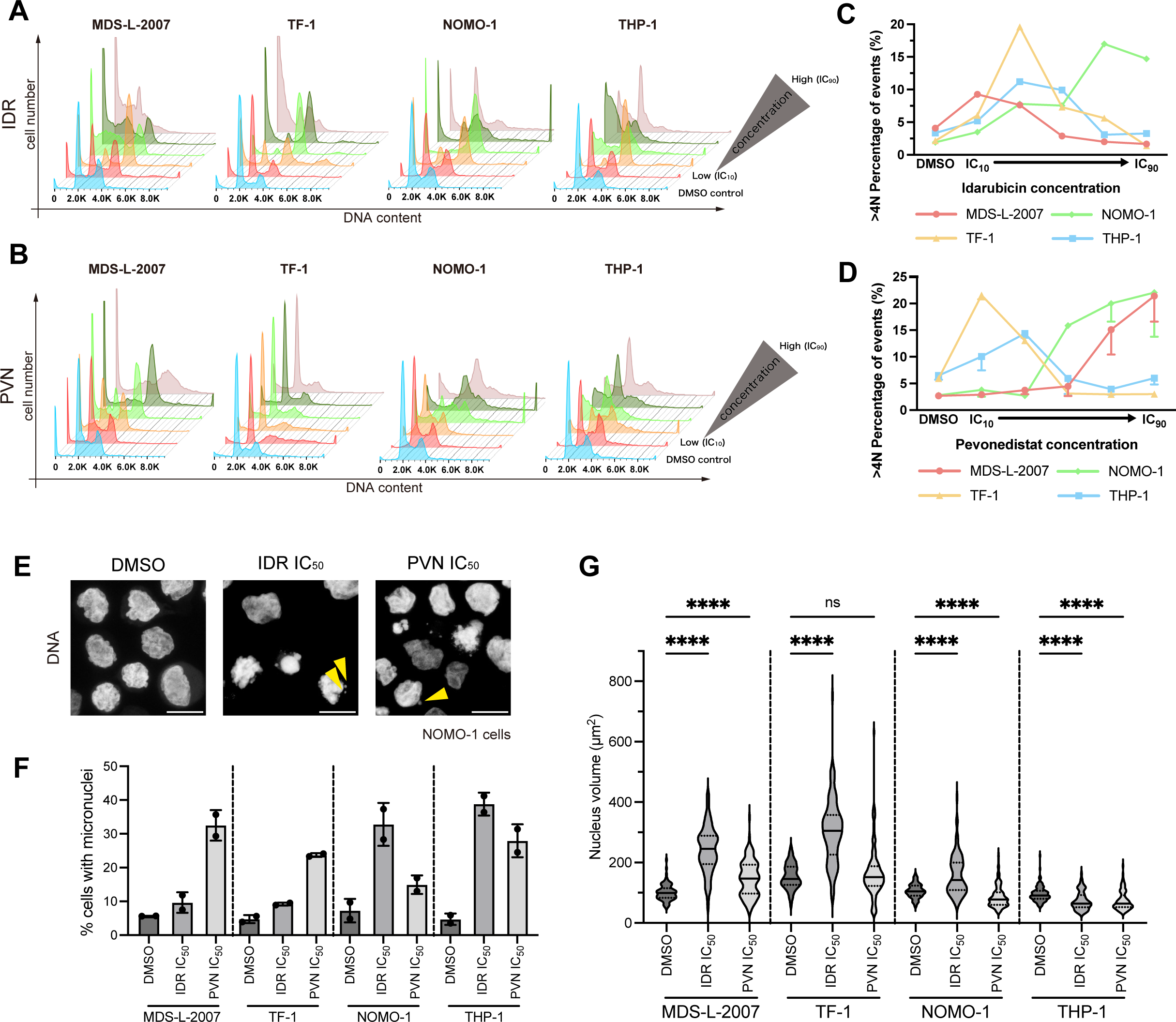
IDR and PVN perturb proper cell cycle progression. **(A) and (B**) Histograms of DNA contents of each AML cell in the presence of various concentrations of drugs. More than 5000 cells were detected. In each panel, the histogram displayed in blue represents the DMSO control. The red histogram, immediately behind the blue histogram, represents approximately the IC_10_ concentration. The histograms further back represent higher concentrations, with the bronze-colored histogram at the very back representing approximately the IC_90_ concentration. Each histogram was created after smoothing processing. (A) and (B) represent IDR and PVN treatment, respectively. In the case of TF-1 cells treated with PVN, the viability only decreased up to 50%, but the histograms towards the back still represent higher concentration conditions. (**C**) The percentage of >4n aneuploid cells in each AML cell line based on the data shown in (A), defining >4n as DNA content >5.0. (**D**) The percentage of >4n aneuploid cells in each AML cell line based on the data shown in (B), defining >4n as DNA content >5.0. n>3. Data are shown as mean - SD. (**E**) Representative images of micronuclei induced by 48 hours treatment with IDR (10 nM) and PVN (40 nM). Scale bar: 10 μm. Arrowheads: micronuclei. (**F**) The percentage of cells with micronuclei in each AML cell line treated with IDR and PVN at approximately IC_50_ concentrations for 48 hours. IC_50_ concentrations was determined based on the data shown in Figure S3D-E. n = 2 independent experiments, over 50 cells for each condition. To exclude nuclear vesicles from dead cells, dead cells with ruptured cell membranes were excluded from the measurement. Data are represented as mean ± SD. (**G**) Violin plot of the nuclear size of each cell at the IC_50_ of IDR and PVN. n > 75. To exclude dead cells from quantification, nuclei with a size of 30 μm^2^ or less were excluded from the analysis. Dunn’s multiple comparisons test was performed to obtain adjusted p-values. ns: not significant, ****p < 0.0001.

**Figure S5.**
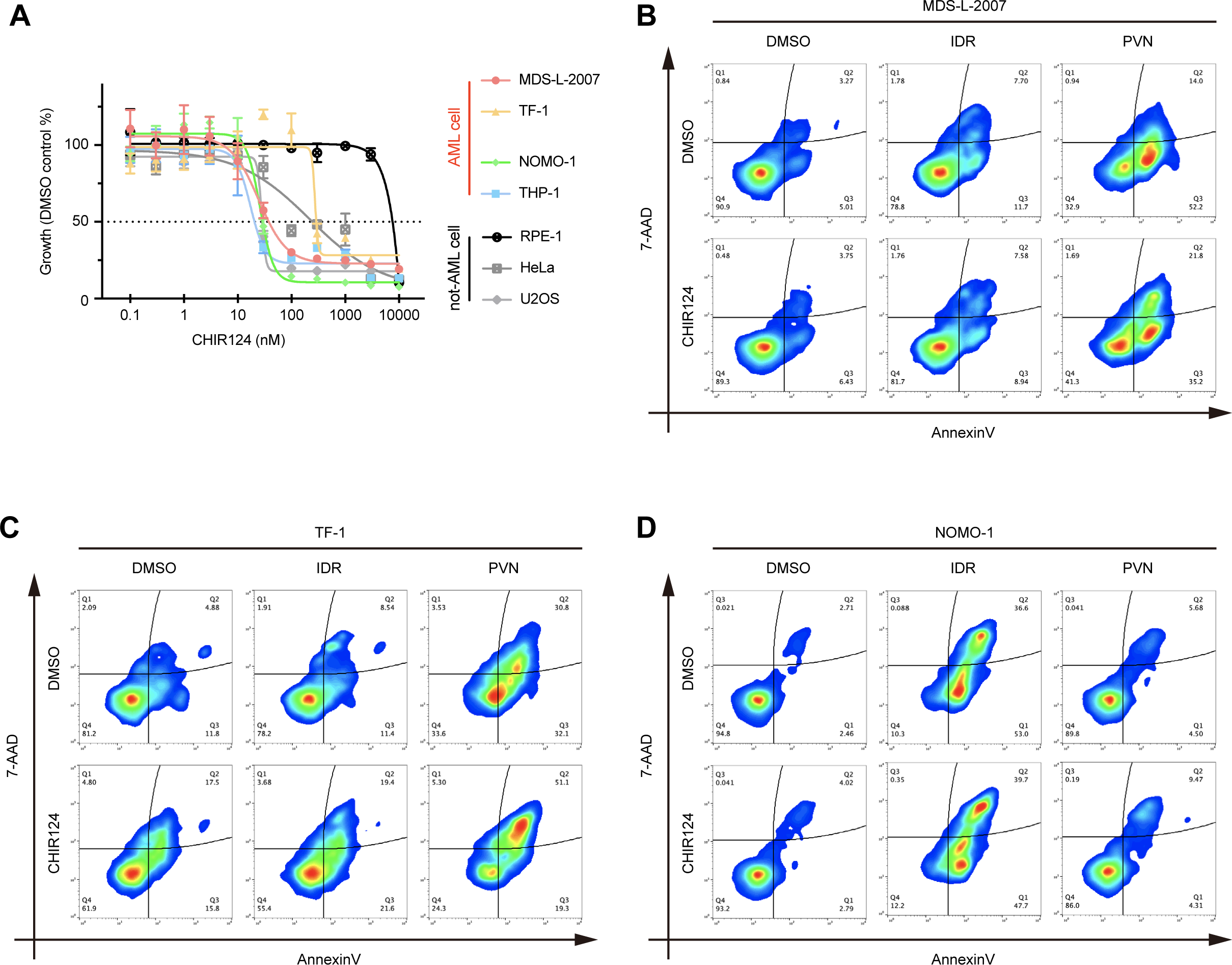
Co-treatment with CHIR124 enhances the cytotoxicity of IDR and PVN. **(A)** Cell survival curves for each cell line were determined by a cell proliferation assay after 48 hours of treatment with the Chk1 inhibitor CHIR124. n>3. Data are shown as mean ± SD. (**B-D**) Heatmap of apoptosis status measured by flow cytometry in AML cells treated with IDR and PVN alone and in combination with CHIR124. CHIR124 was used at a concentration that does not affect cell toxicity over 48 hours. More than 5000 cells were detected. The heatmap was divided into four quadrants (Q1-Q4), and the cells in each quadrant were displayed as percentiles up to 100. Q1 (non-apoptotic cell death: − AnnexinV/+7-AAD), Q2 (late apoptosis/Dead: +AnnexinV/+7-AAD), Q3 (early apoptosis.: +AnnexinV/−7-AAD), and Q4 (living: −AnnexinV/−7-AAD). (B) represents MDS-L-2007 cells. (C) represents TF-1 cells. (D) represents NOMO-1 cells.

